# Ebola virus is evolving but not changing: no evidence for functional change in EBOV from 1976 to the 2014 outbreak

**DOI:** 10.1101/014480

**Authors:** Abayomi S Olabode, Xiaowei Jiang, David L Robertson, Simon C Lovell

## Abstract

The Ebola epidemic is having a devastating impact in West Africa. Sequencing of Ebola viruses from infected individuals has revealed extensive genetic variation, leading to speculation that the virus may be adapting to the human host and accounting for the scale of the 2014 outbreak. We show that so far there is no evidence for adaptation of EBOV to humans. We analyze the putatively functional changes associated with the current and previous Ebola outbreaks, and find no significant molecular changes. Observed amino acid replacements have minimal effect on protein structure, being neither stabilizing nor destabilizing. Replacements are not found in regions of the proteins associated with known functions and tend to occur in disordered regions. This observation indicates that the difference between the current and previous outbreaks is not due to the observed evolutionary change of the virus. Instead, epidemiological factors must be responsible for the unprecedented spread of EBOV.

## Intoduction

The current Ebola epidemic in West Africa is characterized by an unprecedented number of infections, and has resulted in over 8,000 fatalities to date. A recent study using whole-genome sequencing methods ^1^ identified high levels of variation in the Ebola virus (EBOV), much of it unique to the 2014 outbreak. Specifically, 341 substitutions were found of which 35 are non-synonymous, *i.e*., amino acid altering. These changes, coupled with those that occurred prior to the current outbreak, indicate that the virus is evolving rapidly within humans. This presence of the non-synonymous substitutions in the EBOV genomes has led to concern that functional adaptation of the virus to the human host has already occurred ^1,2^, accounting for the unusual scale and severity of the current outbreak.

According to the neutral theory of molecular evolution, mutations may be subject to positive selection if they are beneficial, purifying selection if they are deleterious, or may evolve neutrally if the functional effect is neutral or nearly-neutral ^3,4^. Gire et al suggest that the amino acid replacements observed in the 2014-15 EBOV population are due to incomplete purifying selection ^1^. If this is the case, we would expect replacements to be deleterious and their presence in the population to be due to selection having insufficient time to remove them. By contrast, positive selection may arise if any replacement increases viral fitness. This may be in the form of replacements that stabilize the protein structure, changes that have a beneficial effect on molecular function, or that permit the virus to escape the immune system. Neutral evolution would occur if replacements have minimal effect on fitness. We expect such replacements would be found predominantly in regions of the proteins that are not associated with defined functions, (i.e., away from active sites and interaction sites with other molecules), have minimal effect on protein structure, either positive or negative, and not be in sites subject to selection from the immune system. Neutral evolution is the null hypothesis, and should be assumed in the absence of evidence for either positive or purifying selection.

We use the available sequence data to investigate whether there are deleterious, functional or adaptive change in the EBOV genome. EBOV is a negative stranded RNA virus with a genome comprising seven genes. One of these genes (GP) gives rise to two protein products via transcriptional editing ^5^: a membrane-bound glycoprotein and a secreted protein. Additional functional diversity arises from vp40 existing in a number of conformational forms, which have different functions^6^. Protein structural data are available for at least part of six of the seven EBOV proteins, including three conformational forms for vp40^6-12^. These data allow us to investigate the structural and functional effects of amino-acid replacements in the virus.

## Results

We first investigated whether there is evidence for positive selection acting in any of the seven EBOV genes, using all of the available data (1976 to present). We calculated the ratio between non-synonymous (dN) changes and synonymous changes (dS, non-amino acid altering). A dN/dS ratio close to 1 indicates lack of selection, <1 purifying selection, and >1 that positive selection (adaptive evolution) has taken place. For all seven genes we find few sites with evidence for positive selection (figure 1 and Suppl Table 1). However, except for one case, sites where dN/dS>1 have very low estimates of dS, suggesting that dN/dS ratios are unreliable in this context. For the exception, codon 430 in GP, the single-likelihood ancestor counting (SLAC) model estimates dN/dS to be 4.4. This replacement is not specific to the 2014 outbreak and in a region of the protein predicted to be intrinsically disordered. Disordered regions are known to be permissive for residue changes because they do not have well-defined structures and so are relatively unconstrained^13^. There are also sites with dN but no dS values such that the dN/dS ratio cannot be calculated. As the dN/dS ratio is a relatively conservative method for detecting functional change we computationally characterized all individual amino acid replacements.

**Figure 1.**
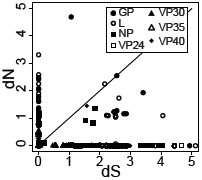
Number of synonymous (dS) and non-synonymous (dN) changes per site. Changes as calculated by the Single-likelihood ancestor counting (SLAC) model ^31^. The line indicates dN=dS.

For the amino acid replacements that fall within a region of known protein structure (figure 2) we assessed the likely functional effect of the substitution. To do this we used phylogenetic methods to reconstruct the most likely sequence of the most recent common ancestor and so the likely evolutionary trajectory. Firstly, we assess the goodness-of-fit of side chains in replacements in all proteins ^14^. We find that all residue changes can be accommodated in the relevant protein structure in a low energy conformation (“rotamer”) with no substantial van der Waals overlaps (figure 3A). Secondly we used an empirical potential to predict the likely structural effect of a residue change ^15^. For all replacements that can be assessed the ΔΔG of folding associated with the amino-acid replacement has a small magnitude compared to the background distribution (figure 3B) indicating that the likely effect on protein structure is minimal. We conclude that all replacements are compatible with their protein structure, and are unlikely to either increase or decrease protein stability.

**Figure 2.**
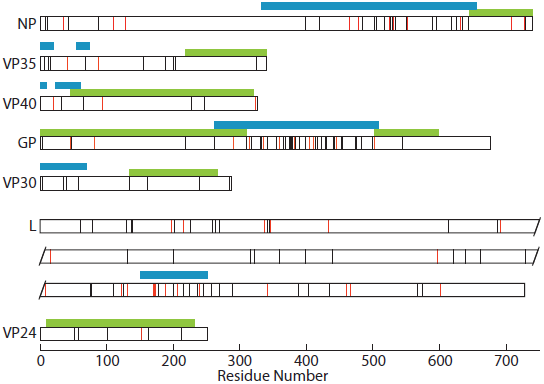
The distribution of amino-acid replacements in the protein sequences. Proteins are shown as white horizontal bars, with replacements indicated by vertical lines. Replacements specific to the 2014 outbreak ^1^ are shown in red. Regions for which the protein structure has been determined are indicated in green, and those regions predicted to be disordered by Disopred ^34^ longer than 10 residues are indicated in blue. The polymerase (L) is broken across three lines for convenient representation.

**Figure 3.**
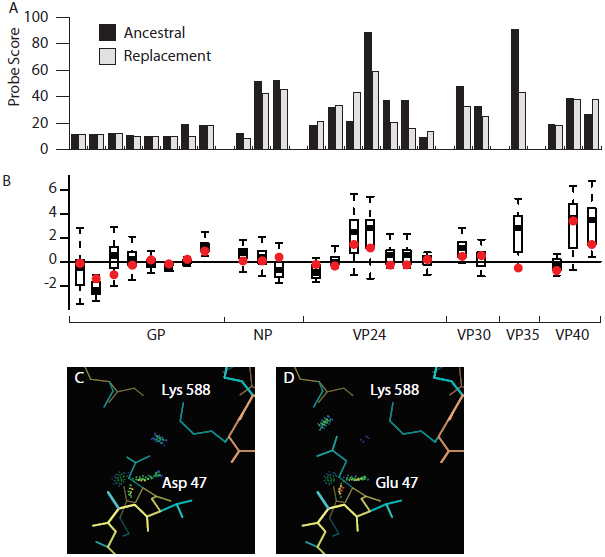
Analysis of amino-acid replacements in the context of protein structure. A. Probe score ^14^ (arbitrary units Å ^2^/x 1000) for ancestral and residues and replacements. Van der Waals overlaps, if present, would result in large negative scores, whereas favorable interactions result in positive scores. B. Change in energy (ΔΔG) for all amino acid replacements found in regions of known protein structure, as predicted using a statistical potential. Each boxplot represents a distribution of energy changes to all 19 other residue types at positions where a non-synonymous substitution has been observed. The ΔΔG of the observed substitution is indicated in red. C. Residue 47 of GP. GP1 is indicated by yellow main chain atoms and GP2 by orange. Van der Waals interactions between the side chain and surrounding atoms are shown by all-atom contact dots ^14^; favorable interactions are colored blue, green and yellow, no unfavorable interactions are found. D. The replacement side chain, modeled in the tt0 rotamer ^30^. Both the ancestral and replacement side chains are negatively charged, and are in close proximity to the positively charged lysine 588 of GP2.

For the GP, VP30 and VP40 proteins, the available protein structures allow identification of protein-protein interaction interfaces, *i.e*., those regions of amino acid sequence responsible for virus to virus and virus to host binding. In total 247 interface residues were identified (from all three proteins, including multiple VP40 conformation forms^6^). Overwhelmingly, replacements in EBOV sequences, whether from the 2014 or earlier outbreaks, are found at sites that do not comprise interaction interfaces. The single exceptions is a replacement of aspartate 47 in GP1 to glutamate, which is not specific to the 2014 outbreak ^1^. Aspartate 47 makes an intra-chain salt bridge with lysine 588 in GP2. Structural modeling of this aspartate to glutamate replacement in GP1 indicates that there is a low energy conformation for glutamate that is able to make the same interaction with no van der Waals overlaps, suggesting this is a relatively minor change (figure 3C). Note, the crystallographic temperature factors are high for this part of the protein structure (many are >100), indicating that the exact positioning of atoms is uncertain. Overall, we conclude that there is no evidence the identified protein interaction interfaces are being disrupted or otherwise altered by the observed amino acid replacements.

As structures are not available for all regions we next investigated the similarity of properties associated with the replacement residues. We find all are conservative for both hydropathy and volume, indicated by the small magnitude of the changes relative to the background distribution (see figure 3 supplement). Larger magnitude alterations are predominantly found in regions of the proteins that are predicted to be intrinsically disordered which are permissive for residue changes. These changes are therefore unlikely to affect structure or function.

Interestingly, half of the amino-acid replacements (88/177) are in regions of proteins predicted to be intrinsically disordered, despite these regions constituting only 27% of the protein sequence (figure 2). In the GP protein the central disordered section corresponds to a mucin-like protein which is highly glycosylated ^5^. Within disordered sequences such as mucin the general character of the amino acid residue is important for function, but, glycosylation sites aside, specific interactions are not made, allowing a range of different amino acids to contribute to the same functional role ^16^.

Disordered regions have been implicated in the formation of new protein interactions ^17^ and may permit viruses to explore novel host perturbations via “sticky” interactions with host proteins ^13^, although predicted disorder is similar for all EBOV outbreaks (figure 4).

**Figure 4.**
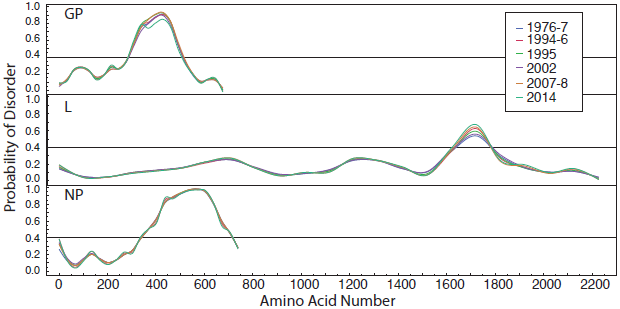
Prediction of disorder for GP, L and NP for sequences from each outbreak. Predictions were made with IUPred ^35^. The recommended cut off (0.4) for considering protein regions disordered is indicated.

A number of experimental studies have identified specific residues that are important for various functions. These include 18 residues in GP that are important for viral entry ^18-20^, four residues in VP30 that are required for nucleocapsid incorporation ^8^, 23 residues in VP24 that are implicated in a range of protein-protein interactions ^11^, four residues in VP40 that make direct contact with RNA ^7^ and a further two residues that are essential for budding ^6^, and three residues in VP35 that are required for binding RNA ^10^. None of these residues are replaced in any of the EBOV lineages sequenced to date, indicating that none of these functions are likely to have been altered, either positively or negatively.

## Discussion

Collectively, the structural and functional data indicate that the observed amino acid replacements are not found in regions of the protein that directly contribute to known functions, and, with the exception of GP aspartate 47, are not in any identified binding interface. None are likely to either stabilize or destabilize the protein structure. The non-synonymous changes are physicochemically conservative, and predominantly cluster in regions predicted to be intrinsically disordered and comparing predictions of these disordered regions for the different EBOV outbreaks indicates this is not changing substantially.

With the exception of a small number of low-frequency frame-shifting intrahost single nucleotide variants, we find no evidence for deleterious mutations that would have been consistent with incomplete purifying selection, as postulated by Gire et al. ^1^. We find no evidence of changes that are likely to be adaptive. This result is corroborated by analysis from a phylogenetic perspective^21^ that similarly shows no evidence of positive selection in the 2014 outbreak. In addition we find no evidence of adaptive change in any of the EBOV sequences from past outbreaks. Although it is not possible to rule out functional change in the intergenic regions, we conclude that none of the non-synonymous substitutions observed to date are likely to affect protein structure or function in any way, be it positive or negative. Thus the null hypothesis of neutral evolution cannot be rejected, and is the most reasonable explanation of the observed sequence diversity.

The lack of functional distinction between the current and previous Ebola outbreaks emphasizes the importance of human-centric epidemiological factors over the molecular biology and evolution of the virus. The main factor that differs between the 2014 outbreak and those that have occurred previously is the establishment of infections in relatively densely populated areas compared with previous outbreaks, coupled with poor health facilities ^22^. Human population growth and progressive urbanization have created efficient pathways for viral transmission ^23^ despite a comparably low basic reproductive number, R_0_ ^22^.

Reconstructing the likely genomes of the most recent common ancestor of each outbreak demonstrates the high degree of similarity of the progenitor virus of each outbreak. This similarity points to a stable viral population in the animal reservoir, which is most likely to be fruit bats ^23^. The zoonotic nature of EBOV and functional stability of the virus suggest that future transmission of similarly virulent potential are highly likely. Given the dense and highly connected nature of the human population, identification of the animal reservoir, surveillance and early intervention will be the key to prevention. The relative functional stability of EBOV suggests that intervention strategies such as with drugs or vaccinations may be more successful than for other RNA viruses.

Strategies for containing EBOV in West Africa ^24^ have been suggested, but are predicated on lack of adaptation of the virus. While evolution of a change in mode of transmission of EBOV is extremely unlikely, due to the highly specific and intricate nature by which viruses interact with their hosts.

Of more concern is the possibility that the R_0_, the number of transmissions per infection, increases. Due to the unusually deadly nature of EBOV, a milder virus with even a substantial drop in virulence would still frequently result in deadly infections on an epidemic, let alone pandemic, scale. Despite observing no functional change between 1976 and mid-2014, future functional change is possible, emphasizing the need for continued monitoring of viral evolution.

## Methods

Ebola sequences were obtained from GenBank, see Gire et al ^1^ for accession numbers. Protein structural data was as follows: NP, pdb code 4QAZ ^12^; VP35, pdb code 3FKE ^10^; GP, pdb code 3CSY ^9^; VP30, pdb code 2I8B ^8^; VP24 pdb code 3VNE ^11^. For VP40, multiple conformations are available, and all were assessed: pdb codes 4LDD, 4LDM, 4LDB ^6^ and 1H2C ^7^.

The sequences were aligned and phylogenetic trees were estimated using the WAG substitution model ^25^, implemented in RAXML ^26^. Ancestral sequences of EBOV proteins was reconstructed using maximum likelihood, implemented by FastML ^27^. Using ancestral reconstruction, the evolutionary pathway for every EBOV sequence in our data set was traced to the last common ancestor, and the sequence of every internal node was compared with that of its ancestor.

The energy change for all amino acid replacements that fall within regions of known structure was predicted using an empirical force field as implemented in FoldX software (version 3 Beta 6). Mutant structures were generated using the “build model” function in FoldX by mutating the native residue in the wild type with the 19 other possible amino acid residues for each position where a non-synonymous mutation was observed. Where the sequence of the native protein differed from that of the crystal structure, the structure was predicted using Modeller ^28^.

Goodness-of-fit of replacement residues in the context of the protein structure was calculated using Probe ^14^ after addition of hydrogen atoms with Reduce ^29^. Probe was also used to identify residues in interaction interfaces. In each case all low energy conformations (“rotamers”) ^30^ were assessed, and the rotamer with the best Probe score used. For replacement valine 325 to isoleucine in VP35, we use the conformation of χ_1_=-45° χ_2_=-50°, which is not the bottom of the energy well, but is nevertheless a favorable side chain conformation.

The SLAC method as implemented by the Datamonkey webserver ^31^ (http://www.datamonkey.org) were used to estimate dN (nonsynonymous) and dS (synonymous) rates for each protein. This method estimates the dN and dS rates for each codon site and compare the observed rates with null expectations based on the used nucleotide substitution model.

Residue volumes were taken from the Zamyatnin ^32^ and hydropathy scales from Kyte and Doolittle, ^33^. Disordered regions of proteins were predicted by DISOPRED ^34^ and lUPred ^35^.

## Acknowledgments

We thank Ryan Ames and Shaun Kandathil for scientific discussion. XJ was supported by MRC (G1001806/1) and Wellcome Trust (097820/Z/11/B) funding to DLR and an award from the Issac Newton Trust/Wellcome Trust ISSF to John Welch at the University of Cambridge.

References

**Figure 3 Supplement.**
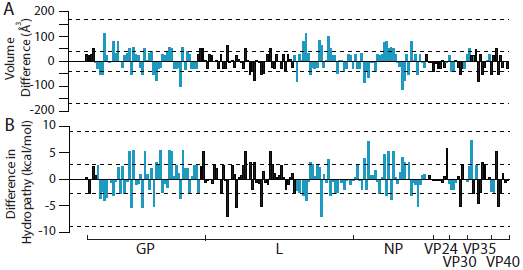
Summary of changes of physicochemical properties arising from all amino acid replacements corresponding to non-synonymous substitutions occurring in EBOV between 1976 and 2014. (A) Volume change for each amino acid residue replacement. Upper and lower bounds, as well as the upper and lower quartiles are indicated by dashed lines. Replacements in regions that are predicted to be disordered are shown in blue, all others in black. (B) Change in hydropathy for each amino acid residue replacement. Upper and lower bounds, as well as the upper and lower quartiles are indicated by dashed lines. Replacements in regions that are predicted to be disordered are shown in blue, all others in black.

## References

1 Gire, S. K. et al. Genomic surveillance elucidates Ebola virus origin and transmission during the 2014 outbreak. Science 345, 1369–1372, doi: 10.1126/science.1259657 (2014).

2 Basler, C. F. Portrait of a Killer: Genome of the 2014 EBOV Outbreak Strain. Cell host & microbe 16, 419–421, doi:10.1016/j.chom.2014.09.012 (2014).

3 Kimura, M. The neutral theory of molecular evolution. (1983).

4 Ohta, T. Slightly deleterious mutant substitutions in evolution. Nature 246, 96–98 (1973).

5 Sanchez, A., Trappier, S. G., Mahy, B. W., Peters, C. J. & Nichol, S. T. The virion glycoproteins of Ebola viruses are encoded in two reading frames and are expressed through transcriptional editing. Proceedings of the National Academy of Sciences of the United States of America 93, 3602–3607 (1996).

6 Bornholdt, Z. A. et al. Structural rearrangement of ebola virus VP40 begets multiple functions in the virus life cycle. Cell 154, 763–774, doi: 10.1016/j.cell.2013.07.015 (2013).

7 Gomis-Rüth, F. X. et al. The matrix protein VP40 from Ebola virus octamerizes into pore-like structures with specific RNA binding properties. Structure (*London, England*: 1993) 11, 423–433 (2003).

8 Hartlieb, B., Muziol, T., Weissenhorn, W. & Becker, S. Crystal structure of the C-terminal domain of Ebola virus VP30 reveals a role in transcription and nucleocapsid association. Proceedings of the National Academy of Sciences of the United States of America 104, 624–629, doi:10.1073/pnas.0606730104 (2007).

9 Lee, J. E. et al. Structure of the Ebola virus glycoprotein bound to an antibody from a human survivor. Nature 454, 177–182, doi: 10.1038/nature07082 (2008).

10 Leung, D. W. et al. Structure of the Ebola VP35 interferon inhibitory domain. Proceedings of the National Academy of Sciences of the United States of America 106, 411–416, doi:10.1073/pnas.0807854106 (2009).

11 Zhang, A. P. P. et al. The ebola virus interferon antagonist VP24 directly binds STAT1 and has a novel, pyramidal fold. PLoS pathogens 8, e1002550, doi:10.1371/journal.ppat.1002550 (2012).

12 Dziubanska, P. J., Derewenda, U., Ellena, J. F., Engel, D. A. & Derewenda, Z. S. The structure of the C-terminal domain of the Zaire ebolavirus nucleoprotein. Acta crystallographica Section D, Biological crystallography 70, 2420–2429, doi:10.1107/S1399004714014710 (2014).

13 Xue, B. & Uversky, V. N. Intrinsic disorder in proteins involved in the innate antiviral immunity: another flexible side of a molecular arms race. J Mol Biol 426, 1322–1350, doi:10.1016/j.jmb.2013.10.030 (2014).

14 Word, J. M. et al. Visualizing and quantifying molecular goodness-of-fit: small-probe contact dots with explicit hydrogen atoms. J Mol Biol 285, 1711–1733, doi: 10.1006/jmbi.1998.2400 (1999).

15 Guerois, R., Nielsen, J. E. & Serrano, L. Predicting changes in the stability of proteins and protein complexes: a study of more than 1000 mutations. Journal of Molecular Biology 320, 369–387, doi: 10.1016/S0022-2836(02)00442-4 (2002).

16 Nishikawa, I. et al. Computational prediction of O-linked glycosylation sites that preferentially map on intrinsically disordered regions of extracellular proteins. International journal of molecular sciences 11, 4991–5008, doi:10.3390/ijms11124991 (2010).

17 Meszaros, B., Simon, I. & Dosztanyi, Z. Prediction of protein binding regions in disordered proteins. PLoS computational biology 5, e1000376, doi:10.1371/journal.pcbi.1000376 (2009).

18 Brindley, M. A. et al. Ebola virus glycoprotein 1: identification of residues important for binding and postbinding events. Journal of virology 81, 7702–7709, doi:10.1128/JVI.02433-06 (2007).

19 Manicassamy, B., Wang, J., Jiang, H. & Rong, L. Comprehensive analysis of ebola virus GP1 in viral entry. Journal of virology 79, 4793–4805, doi:10.1128/JVI.79.8.4793-4805.2005 (2005).

20 Mpanju, O. M., Towner, J.S., Dover, J. E., Nichol, S. T. & Wilson, C.A. Identification of two amino acid residues on Ebola virus glycoprotein 1 critical for cell entry. Virus research 121, 205–214, doi: 10.1016/j.virusres.2006.06.002 (2006).

21 Spielman, S. J., Meyer, A.G. & Wilke, C. O. Increased evolutionary rate in the 2014 West African Ebola outbreak is due to transient polymorphism and not positive selection. Bioarxiv doi: http://dx.doi.org/10.1101/011429 (2014).

22 Gatherer, D. The 2014 Ebola virus disease outbreak in West Africa. The Journal of general virology 95, 1619–1624, doi: 10.1099/vir.0.067199-0 (2014).

23 Pigott, D. M. et al. Mapping the zoonotic niche of Ebola virus disease in Africa. eLife 3, e04395, doi:10.7554/eLife.04395 (2014).

24 Pandey, A. et al. Strategies for containing Ebola in West Africa. Science 346, 991–995 (2014).

25 Whelan, S. & Goldman, N. A general empirical model of protein evolution derived from multiple protein families using a maximum-likelihood approach. Mol Biol Evol 18, 691–699 (2001).

26 Stamatakis, A. RAxML version 8: a tool for phylogenetic analysis and post-analysis of large phylogenies. Bioinformatics 30, 1312–1313, doi: 10.1093/bioinformatics/btu033 (2014).

27 Ashkenazy, H. et al. FastML: a web server for probabilistic reconstruction of ancestral sequences. Nucleic acids research 40, W580–584, doi: 10.1093/nar/gks498 (2012).

28 Sali, A. & Blundell, T. L. Comparative protein modelling by satisfaction of spatial restraints. J Mol Biol 234, 779–815, doi: 10.1006/jmbi.1993.1626 (1993).

29 Word, J. M., Lovell, S. C., Richardson, J. S. & Richardson, D. C. Asparagine and glutamine: using hydrogen atom contacts in the choice of side-chain amide orientation. J Mol Biol 285, 1735–1747, doi: 10.1006/jmbi.1998.2401 (1999).

30 Lovell, S. C., Word, J. M., Richardson, J. S. & Richardson, D. C. The penultimate rotamer library. Proteins 40, 389–408 (2000).

31 Kosakovsky Pond, S. L. & Frost, S. D. W. Not so different after all: a comparison of methods for detecting amino acid sites under selection. Molecular biology and evolution 22, 1208–1222, doi: 10.1093/molbev/msi105 (2005).

32 Zamyatnin, A. A. Protein volume in solution. Progress in biophysics and molecular biology 24, 107–123 (1972).

33 Kyte, J. & Doolittle, R. F. A simple method for displaying the hydropathic character of a protein. J Mol Biol 157, 105–132 (1982).

34 Ward, J. J., McGuffin, L. J., Bryson, K., Buxton, B. F. & Jones, D. T. The DISOPRED server for the prediction of protein disorder. Bioinformatics 20, 2138–2139, doi:10.1093/bioinformatics/bth195 (2004).

35 Dosztanyi, Z., Csizmok, V., Tompa, P. & Simon, I. IUPred: web server for the prediction of intrinsically unstructured regions of proteins based on estimated energy content. Bioinformatics 21, 3433–3434, doi: 10.1093/bioinformatics/bti541 (2005).

